# Dominant constraints on the evolution of rhythmic gene expression

**DOI:** 10.1101/2022.06.14.496129

**Authors:** Yang Cheng, Yuhao Chi, Linying Sun, Guang-Zhong Wang

**Author notes:** Corresponding author (W. G-Z.). These authors contributed equally to the paper (Yang Cheng, Yuhao Chi, Linying Sun).

## Abstract

Although the individual transcriptional regulators of the core circadian clock are distinct amongst different organisms, the autoregulatory feedback loops they form are conserved. This unified design principle explains how daily physiological activities oscillate across species. However, whether analogous design principles govern the gene expression output of circadian clocks is unknown. Herein, we performed a comparative analysis of rhythmic gene expression in eight diverse species and captured four common distribution patterns of cycling gene expression across these species. We hypothesized that maintenance of reduced energetic costs constrains the evolution of rhythmic gene expression. Our large-scale computational simulations support this hypothesis by showing that selection against high-energy expenditure completely regenerates all cycling gene patterns. Moreover, we find that the late- and early-phase peaks of rhythmic expression have been subjected to this type of selective pressure. Therefore, energetic costs have constricted the cycling transcriptome throughout evolutionary history.

## Main

The circadian clock is a conserved time-entrained mechanism of living organisms that affects all aspects of physiology^1–4^. The regulatory proteins that make up the core circadian clock are markedly different between species^5, 6^. In mammals, the positive regulators are BMAL1 and CLOCK/NPAS2, which activate the repressors PERs and CRYs, RORs and REV-ERBs, or DBP^1, 7, 8^. These negative regulators subsequently repress their upstream activators. In Drosophila, CLK and CYC form heterodimers to induce either PER and TIM or VRI and PDP1, which repress or activate CLK, respectively^9, 10^. In cyanobacteria, the essential elements are KaiA, KaiB, and KaiC^11, 12^. In Arabidopsis, interconnected circuits joining morning- and evening-specific transcriptional loops contain the proteins CCA1, LHY, and TOC1^13–15^. Regardless of the actual constituents, all molecular clocks ultimately exert temporal and spatial control of transcription through a negative feedback motif^3, 16^. This autoregulatory negative feedback loop oscillates with the 24-hour day and explains the molecular mechanisms controlling circadian adaptation to the environment.

The gene-expression outputs of circadian regulatory networks also vary between different conditions and species. Early microarray-based investigations on cycling transcripts in the liver and SCN of mice found that, while hundreds of genes are rhythmically expressed in the two tissues, the overlap between them is small^17^. This finding is consistent with large-scale comparisons of multiple organs either in mice and baboons^18, 19^. Differential rhythmic expression is also observed between light/dark and dark/dark behavioral paradigms in mice, with more diurnal transcripts being detected under L/D conditions^20^. Likewise, the proportion of the cycling transcriptome decreases in aged tissues^21^, and distinct feeding regimens greatly alter the number of rhythmically-expressed genes^22, 23^. Similar results have been reported in other species such as Drosophila^19, 24, 25^.

In the present study, we hypothesized that conserved patterns of rhythmic gene output organize the circadian transcriptome across species, analogous to what is seen in the regulatory motif of the core circadian clock. We tested this hypothesis by comparing datasets of rhythmic gene expression from eight species to discover four common distribution patterns that govern circadian transcriptional output. We hypothesized that a model in which selection against high energetic cost constrains the evolution of the cycling transcriptome. Our calculation suggests that in yeast the rhythmic expression of most genes is under this constraint, but that value is less than 10% of the rhythmic genes in mice. Our model further indicates that the co-expression of only a few murine genes can reach the selection threshold, highlighting the important role of phase alignment. Both the early- and late-phase peaks characteristic of transcriptional oscillation are subject to this selection. Finally, our computational simulations based on this cost optimization model can fully reproduce all four gene patterns observed. Our results provide insights on unified design principles underlying circadian gene expression across species.

## Results

### Cycling Genes are Distinct Between Species

We sought to determine to what extent the circadian transcriptome is conserved across species. We thus compared cycling transcriptomes from 8 species under different experimental conditions, including human^26, 27^, mouse^18^, *Drosophila melanogaster*^28^, yeast^29^, *Neurospora crassa*^30^, *Arabidopsis thaliana*^31^, *Chlamydomonas reinhardtii* (green algae)^32^ and *Synechococus elongates*^33^ (Figure 1A). These 8 species cover the phylogeny represented by model systems used in circadian biology, and the data cover a wide range of tissues and conditions (Figure 1A, Data Table S1). We noted that those datasets were from both RNA-seq and microarray profiling, due to the limitation of data availability. However, these 14 cycling datasets allowed us to reveal the essential characteristics of oscillating gene, as both techniques have been proved that they can provide effective biological signals. What is more important, the conserved patterns of rhythmic genes that we aim to search for should be detected regardless of experimental conditions, organs and sequencing platforms.

**Figure 1.**
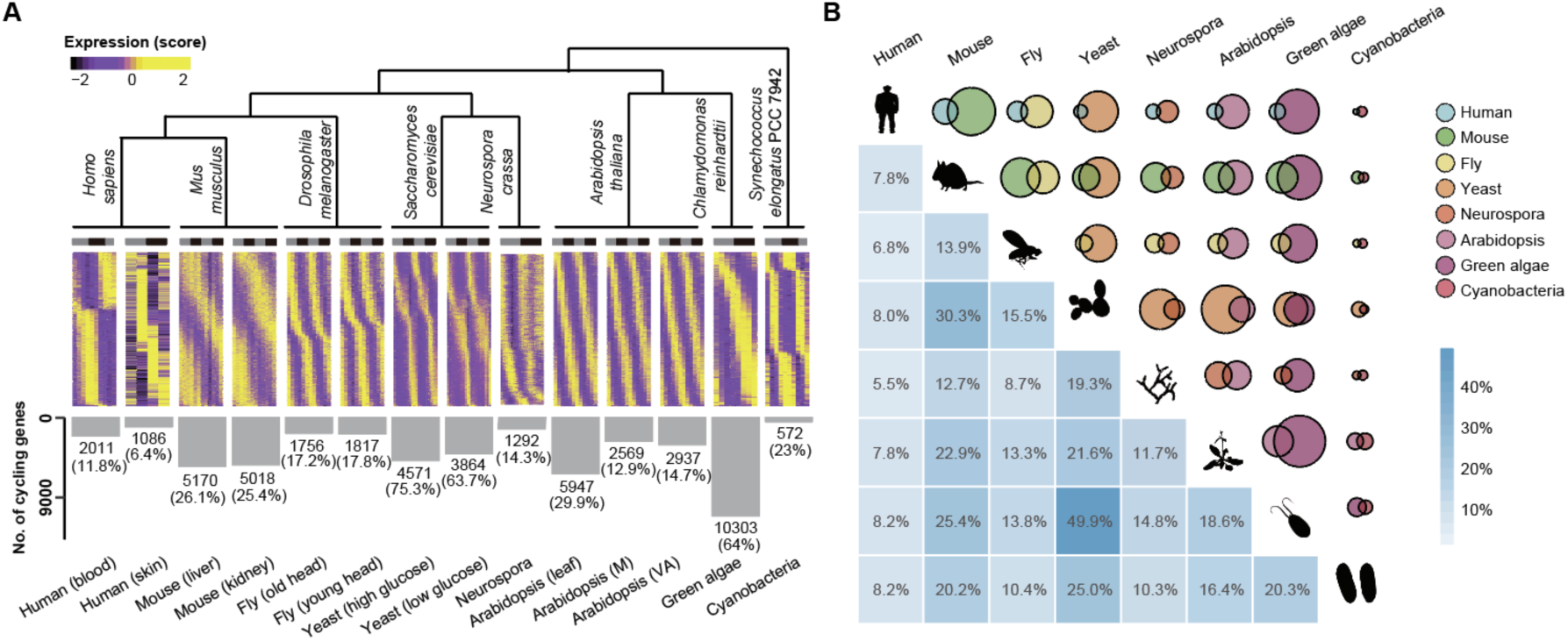
Cycling gene expression in 8 species. (A) Heatmaps of rhythmic gene expression in eight species, including *Homo sapiens*, *Mus musculus*, *Drosophila melanogaster*, *Saccharomyces cerevisiae*, *Neurospora crassa*, *Arabidopsis thaliana*, *Chlamydomonas reinhardtii* and *Synechococus elongates*. The number and proportion of identified cycling genes in each transcriptome are indicated near the bottom of the panel. Cycling genes were identified by MetaCycle with adjusted p values < 0.05 after a Benjamini-Hochberg procedure. (B) Venn diagrams displaying the intersection of cycling genes between any two species. Different colors represent different species, while the size of the circle indicates the number of cycling genes identified. The proportion of the overlaps are indicated below, with higher percentages highlighted by a darker color. Average values were used for species with two or more associated tissues/conditions.

In total, 193,492 transcripts were searched for an oscillating expression signature via the “MetaCycle” package^34^, with Benjamini-Hochberg multiple-comparison corrections of q < 0.05 being applied (Figure 1A, Data Table S2). We found that the absolute number and relative proportions of cycling transcripts varies to a great extent across species (Figure 1A). In green algae, 10,303 genes were detected as rhythmically expressed, which is 64% of all transcribed genes, while in human skin tissue, only 1,086 (6.39%) genes were determined to be cycling genes. We also found that yeast displayed the highest proportion of cycling transcripts (under high glucose conditions), which covers as much as 75.32% (4,571/6,069) of its transcriptome. In mammals including humans and mice, approximately 6% - 26% of transcripts were rhythmically expressed. Unicellular eukaryotic organisms (*Chlamydomonas reinhardtii*, yeast, *Neurospora crassa*, and *Synechococcus elongates*) possess a proportionally larger cycling transcriptome than multicellular organisms (48.05% versus 18.02%) (Figure 1A), which indicates that the genomes of unicellular organisms more readily anticipate periodicity within the environment.

Next, we asked what portion of the genes that exhibit rhythmic expression are shared across different species. Previous studies have only performed comparisons that covered a limited number of species or experimental conditions. To better answer this question, pairwise reconstructions of orthologous gene groups between all pairs of species were performed, which covered 8×7 = 56 species pairs and 21,075 ortholog gene groups. Our results show that the proportion of overlapping cycling genes between any two species is small (Figure 1B), with an average of 15.97% (404 cycling genes). The largest number of shared cycling genes occurred between yeast and green algae (1,227 genes) and the smallest number of shared cycling genes was found between human and *Synechococcus elongates* (19 genes). Only 446 cycling genes were common between human and mouse tissues, or 7.8% of the whole cycling transcriptome. In addition, the number of shared cycling genes decreases dramatically as the number of species under comparison increases (Figure 2B). Finally, our analysis including all species suggests that there is no relationship between the proportion of shared cycling genes and their evolutionary distance (Figure S1A-H, p > 0.64 for all the comparisons). These results illustrate that cycling genes rarely overlap between species.

**Figure 2.**
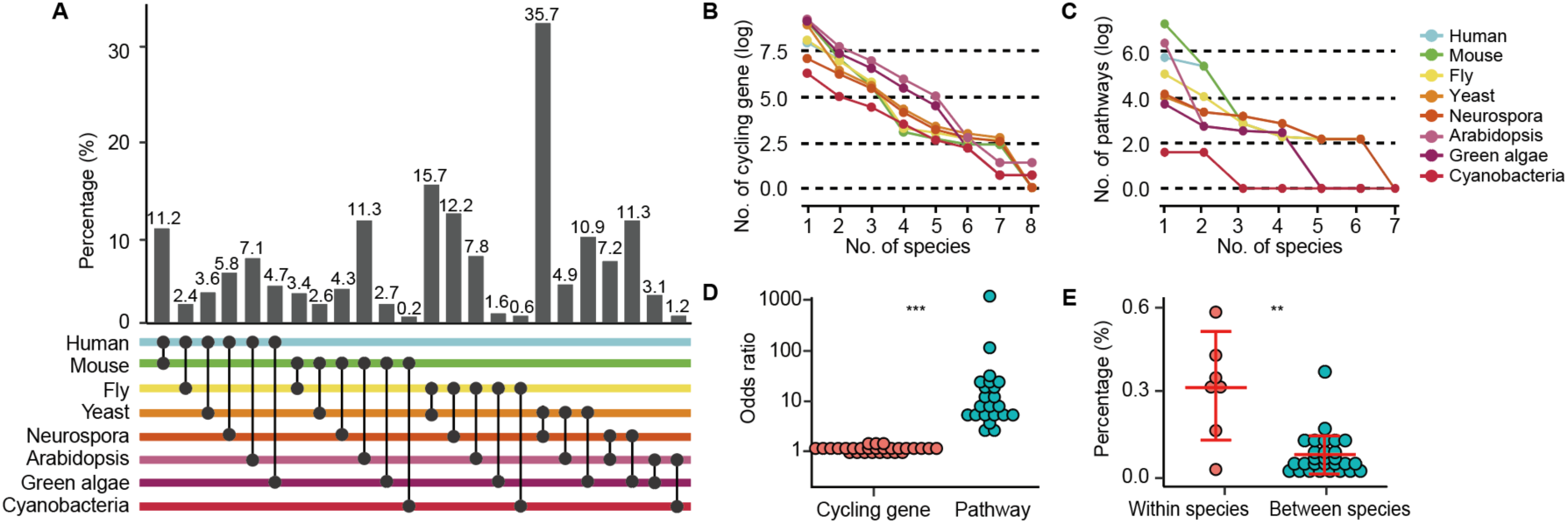
Functional enrichment analysis of cycling genes across species. (A) The proportion of conserved pathways between any species pair. Bar plots represent the proportion of conserved pathways, which were calculated by the number of shared enriched pathways divided by the total number of enriched pathways between the two species. Different species are labeled by distinct colors and detailed comparison information is highlighted below the panel. (B) The trends of shared cycling genes across species. As more species are included, the number of shared cycling genes drops significantly. Lines of different colors represent distinct species and values obtained after logarithmic transformation are shown. (C) The trends of shared functional pathways of cycling genes across species. Only enriched pathways are included in the plot. The number of shared enriched pathways of cycling genes drops significantly as more species are considered. Lines of different colors represent distinct species and values obtained after logarithmic transformation are shown for the number of enriched pathways. (D) Dot plot showing the comparison of shared cycling genes versus their enriched pathways across species. Odds ratio of both shared cycling genes and shared pathways between species pairs are plotted, and statistical significance was determined by a Wilcoxon rank sum test (*** indicates p < 0.001). (E) Dot plot showing the comparison of conserved enriched pathways between and within species. Proportion of conserved enriched pathways is plotted and significance levels were determined by a Wilcoxon rank sum test (** indicates p < 0.01).

### Functional Pathways Enriched for Cycling Genes are Organism-Specific

We next asked whether cycling genes participate in similar biological functions among different species. We therefore explored the enrichment of biological pathways amongst cycling genes by utilizing comprehensive Gene Ontology analysis^35–37^. Indeed, multiple pathways were uncovered to be overrepresented in cycling genes (Data Table S3). For instance, in yeast, 24 biological functions are significantly enriched for cycling genes (high glucose conditions), while in green algae 42 biological pathways are overrepresented. However, our combined analysis suggests that only 8 biological pathways are both overrepresented by the cycling genes in the two organisms, which is only 13.79% of the total number of enriched pathways (Figure 2A). On average, the number of shared enriched biological pathways for yeast and green algae are 7 (10.94%) (Figure 2A). Overall, the pairwise analysis of any two species shows an enrichment of 0 to 35.71% of shared biological pathways for rhythmic gene expression, with a median proportion of 3.98% and an average of 6.13% (Figure 2A). Similar to what was observed at the level of individual genes (Figure 2B), the number of shared pathways decreased as the number of compared species increased and no common enriched pathways were detected across all eight species (Figure 2C). Additionally, evolutionary distance did not correlate with the proportion of shared pathways among these organisms (Figure S2A-H). The above results suggest that biological pathway enrichment is not a conserved feature of rhythmic gene expression across species.

Next, we asked whether the overlap of cycling genes at the pathway level is quantifiably larger than what was observed at the individual level. Rhythmic gene identity was significantly more conserved at the pathway level than at the gene level (median enrichment odds ratio = 5.62 versus 1.05, respectively, *p* = 1.33×10^-5^, Wilcoxon rank sum test, Figure 2D). Similarly, the proportion of conserved pathways is higher than the proportion of shared cycling genes in both mice (p = 5.6×10^-15^, Figure S3A) in Arabidopsis (p = 2.5×10^-6^, Figure S3B). These results show slight conservation of cycling gene functions between species and among different tissues.

Interestingly, it seems that the number of shared biological pathways enriched for cycling genes within one species is higher than that between species (Figure 2E, p = 0.006, Wilcoxon rank sum test). For instance, while there are only 2.62% enriched pathways that are shared between yeast and mouse circadian transcriptomes, 61.11% pathways are found to be enriched in yeast high- and low glucose conditions, and 34.68% pathways are commonly enriched between mouse circadian tissues (liver and kidney). The high proportion of shared circadian pathways within organisms, rather than between organisms, is likely a result of local environmental adaptation by each species.

### Cycling Genes are Highly Expressed and Exhibit a Large Range of Expression

We first considered whether or not genes regulated by the cycling transcriptome display a large expression range compared to non-cycling genes. We calculated the relative amplitude (rAMP) of cycling genes and defined these values as cycling regulation ranges, where rAMP is defined as the ratio between amplitude and baseline expression and obtained from the output of “MetaCycle”. Cycling genes were highly expressed (Figure 3A)^29, 38^ and displayed a larger average regulation range than non-cycling genes in all species considered (Figure 3B). We then tested whether the proportion of cycling genes increases with the relative expression range, as this uptrend was universally observed against expression levels (Figure 3C). By classifying all the cycling genes into five equal groups according to their expression range, we plotted the distribution of cycling genes. In all species, the proportion of cycling genes gradually increases in the large rAMP categories (Figure 3D). For instance, in the mouse liver, only 2.65% of cycling genes are in the bottom 20% of ranked genes, while 42.42% of cycling genes are located in the top 20%. These results hint that highly-expressed cycling genes have a greater flexibility in their expression than non-cycling transcripts and that rhythmicity regulates gene activity at scale. In summary, both expression level and regulation range are evolutionarily conserved traits that control the distribution of cycling genes.

**Figure 3.**
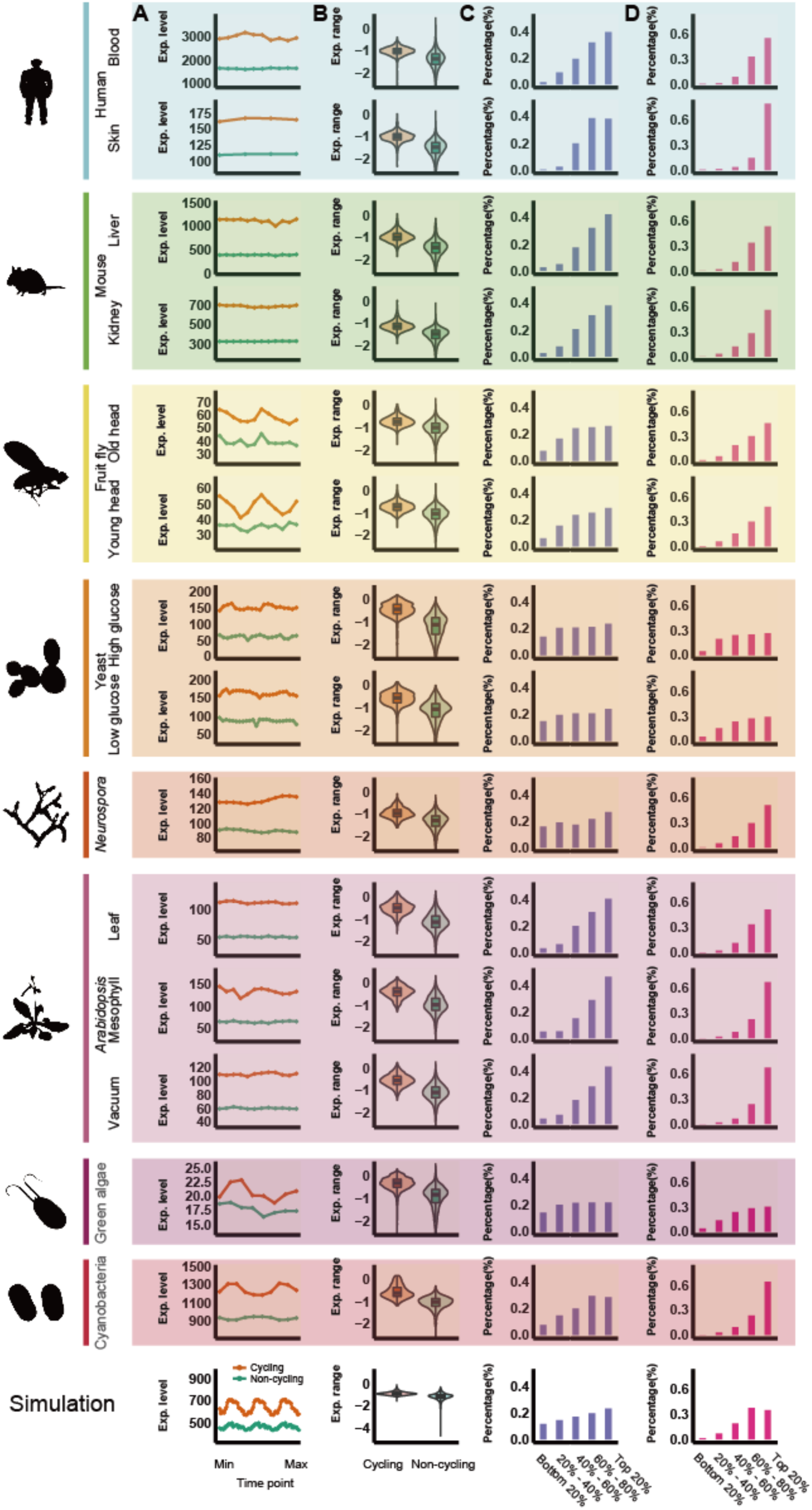
Conserved distribution patterns of cycling genes across species. (A) Mean expression levels of cycling genes versus other genes for each time point. Red lines represent cycling genes while green lines represent non-cycling genes in each plot. p values < 0.005 for all comparisons between the expression of cycling genes with other genes for each time point (total of 244 comparisons), Wilcoxon rank sum test. (B) Violin plots showing comparisons of relative regulation ranges between cycling genes and other non-cycling genes. Red boxes represent cycling genes while green boxes represent non-cycling genes. p < 7.046×10^-190^ in each of the 15 comparisons, Wilcoxon rank sum test. (C-D) Proportions of cycling genes in each category grouped by mean expression level (C) and rAMP. (D). Five groups were calculated, from the bottom 20% to the top 20%. Percentages were calculated as the number of cycling genes in that group divided by the total number of cycling genes identified in the cycling transcriptome of each species. Logarithmic values for both of the parameters are shown.

Do highly expressed genes tend to be detected as cycling genes and thus explain the four observed patterns? There are several reasons that this scenario is unlikely. Firstly, the link between cycling genes and high expression cannot explain the association between cycling genes and large regulation range, as expression level and rAMP are generally not positively correlated (Figure S4). Secondly, if we assumed that all the expressed genes were downregulated, the detected cycling genes would be reduced according to the cycling genes misclassification hypothesis. We surprisingly discovered that the total number of cycling genes in each transcriptome was unchanged if the expression of each gene were downregulated by 50%. Thirdly, we artificially generated 10,000 expressed genes, with 2,500 (1/4 of them) cycling genes included which were evenly distributed along with expression level. It seems that the cycling genes are still evenly distributed in distinct expression groups if we increased the variation of expression of each gene by 20% or 50% (Figure S5). Fourthly, our observed patterns remains if we removed the bottom 20% of expressed genes, as they might contain more “noise” (Figure S6). Similarly, oxidation-reduction cycles^39^ cannot explain those patterns as if oxidation-reduction related genes were removed from the analysis, all the four patterns still can be observed.

### Prediction of Cycling Genes Based on Expression and Relative Regulation Range

As both expression ranges and levels can affect the distribution of cycling genes, we sought to predict whether or not specific genes are rhythmic based on the two parameters. To accomplish this, we first selected gene sets whose rAMP covered >95% of the total cycling genes. We then calculated the number of correctly predicted cycling genes (Figure 4A-N, also see METHOD DETAILS). Using this approach, highly-expressed genes have a higher chance of being selected as a cycling gene. We found that in the mouse liver, up to 62.57% of cycling genes can be correctly predicted by this protocol (3235 of 5,170 cycling genes). In yeast under high-glucose conditions, about 90% of cycling genes can be correctly predicted (4068 of 4,571 cycling genes). The results from 1,000 computational experiments show that the number of correctly predicted cycling genes is significantly higher than expected by chance (p < 2.7×10^-165^ in all the comparisons).

**Figure 4.**
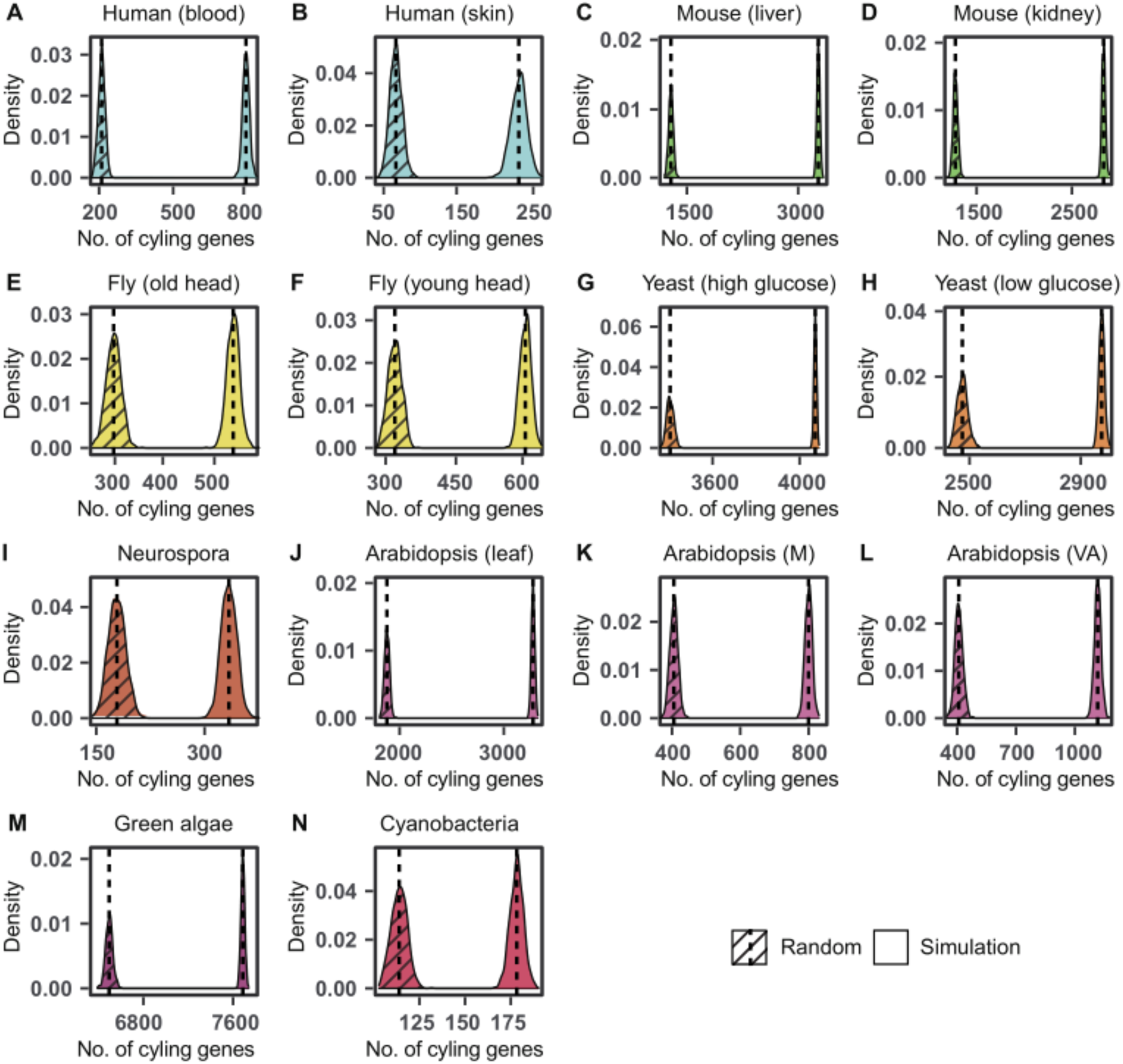
Prediction of cycling genes based on expression level and relative regulation range. Results are plotted for all species (A-N), with controls indicated by slashes. Controls were defined as predictions of cycling genes without considerations for expression levels or regulation ranges. Different species are distinguished by different colors and 1,000 simulations were performed for each distribution curve. Vertical lines represent the means of each distribution peak.

### Energetic Cost Constrains Rhythmic-gene Expression

Provided with the above results, we hypothesized that these patterns could be explained by the pressure of reducing energetic cost during gene expression. Highly-expressed genes require more energy (ATP); reducing energy expenditure associated with their transcription and translation is thus favored. Activating the expression of genes when they are needed and repressing their expression when they are not is an effective strategy to achieve optimization of transcriptional and translational energetics. This strategy benefits the whole transcriptome and is critical for survival during limited nutrition. For population genetics, the selective coefficient is determined by the effective population size^40, 41^ and natural selection is more efficient in species with large effective populations. For the 8 species investigated in this study, the effective population sizes of yeast, *Neurospora crassa*, green algae, and cyanobacteria are much larger than those of humans, mice, fruit flies, and Arabidopsis. The larger the effective population size, the smaller the selective coefficient (s = 4/Ne)^40, 41^, indicating that a small change in the energetic cost of gene expression is sufficient to be detected by natural selection^41–44^. Detailed experimental measurements required for the calculation of energy expenditure, such as the genome-wide measurement of mRNA and protein half-lives, together with decay rates for each gene, have been previously reported^41, 42, 45^. We thus evaluated the selective constraints of energetic cost on the cycling transcriptomes of yeast (under high-glucose conditions) and mice (liver tissue) (Figure 5A).

**Figure 5.**
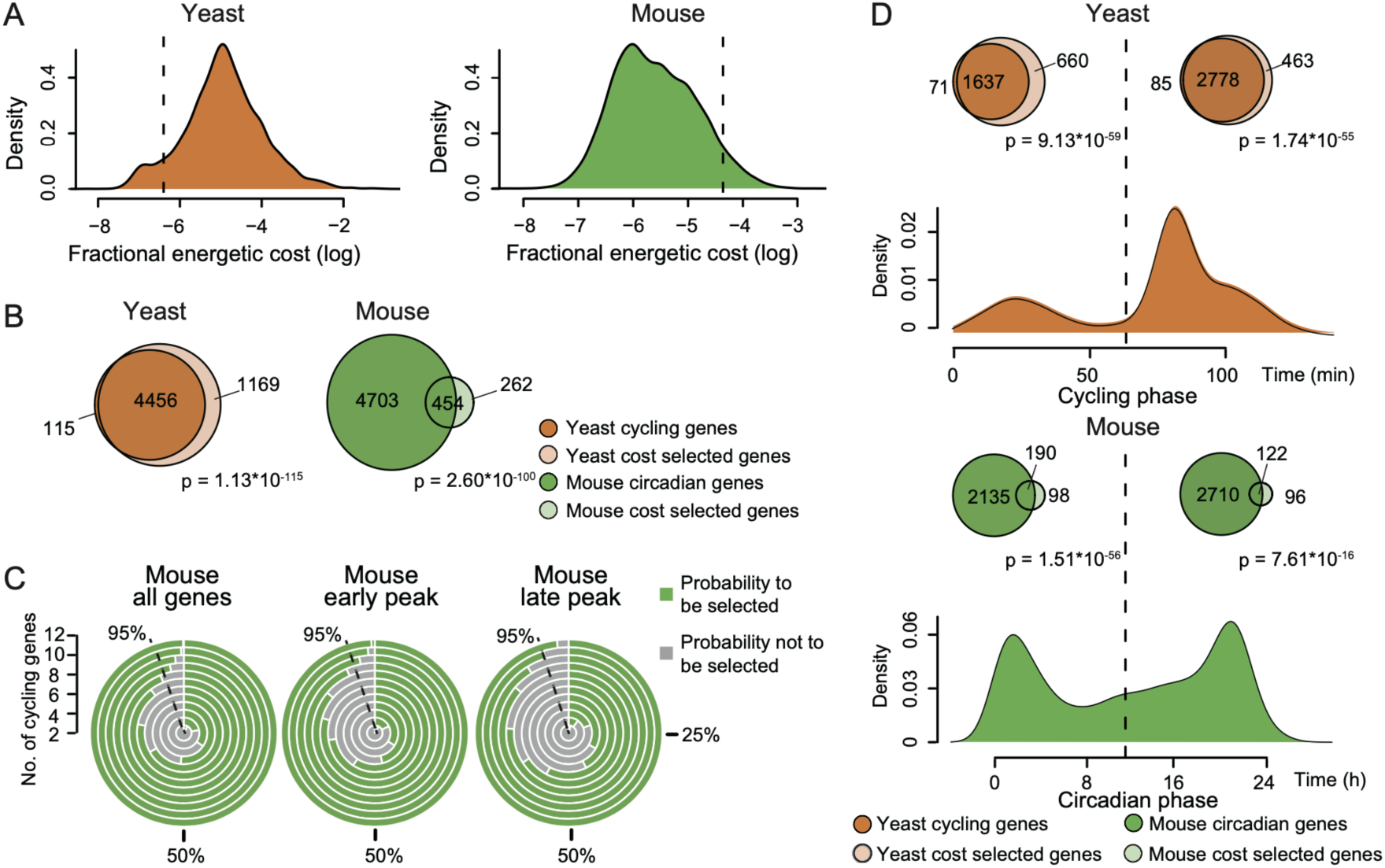
Energy cost constrains the cycling transcriptomes of yeast and mice. (A) Distribution of fractional cost for each gene in the cycling transcriptome of yeast and mice. Vertical lines represent the threshold for selection, which was determined by the effective population size. (B) Venn diagrams illustrate the intersection between cycling genes and genes whose expression passed the cost selection threshold in yeast and mice. Odds ratio = 10.87 and 5.23 for yeast and mice, respectively. (C) Radial bar chart visualization on the selection of co-expressed genes. Equal or greater than 95% chance to be under selection is indicated by a dash line. (D) Enrichment of cost-selected genes in both the early and late circadian expression peaks. Venn diagrams showing the overlaps between cost-selected genes and cycling genes whose phase is located in either the early or late peak. Odds ratio = 8.17 and 9.87 for yeast early and late peaks, respectively; Odds ratio = 7.04 and 3.06 for mouse early and late peaks, respectively. Fisher’s exact tests were used to determine significance levels for each enrichment.

We determined the selective coefficient (1.47×10^-7^) in yeast by the species’ reported effective population size (1.36×10^7^)^41^. Selection pressure is non-negligible if the cost change exceeds this threshold (1.47×10^-7^) during rhythmic expression. For the 4,571 periodically expressed genes, the cost alteration for 4,456 (97.48%) genes is above the selective coefficient during yeast rhythmic expression (high glucose condition) (Figure 5B, Data Table S4). Even for the majority of all 6,068 expressed yeast genes (5,625, 92.70%), dynamic expression has passed the selective coefficient threshold (Figure 5A). Thus, for an organism with a large effective population size, rhythmic-gene expression effectively removes transcripts that are not relevant at any particular time by selecting against the cost of their expression.

As the effective population size of mice is much smaller than that of yeast (72,500 versus 1.36×10^7^)^41, 46^, the selective coefficient for the mouse population is two magnitudes larger than that for the yeast population (1.47×10-7 versus 5.5×10^-5^, respectively). Unsurprisingly then, for the 5,175 cycling genes in mice, the oscillation of only 454 (8.77%) genes is under selection by energetic-cost optimization (Figure 5B). For all 19,621 mouse genes, only 716 (3.65%) passed the selective threshold in the cycling transcriptome (Figure 5A, Data Table S4). Still, cost-selected genes are enriched amongst cycling genes exists in both yeast and mice (OR = 10.87 and 5.23, p < 2.60×10^-100^, respectively, Figure 5B).

The above results suggest that the cyclic expression of individual genes rarely reaches the selective threshold in mice, due to the small effective population size of this species. As cycling genes are often co-expressed in order to meet related demands, we assessed the number of phase-aligned genes that are required to reach the selective coefficient. On average, we found that as few as 9 co-oscillating genes (with similar peak phases) can be detected by selecting for energetic cost with 95% accuracy (Figure 5C). These results suggest that selection pressure applied to the peak phase alignments of small sets of genes is an effective way to remove temporally-unneeded transcripts from the mouse transcriptome.

Generally, the strength of selective pressure on energetic cost positively correlates with gene expression in yeast (r = 0.724, p < 10^-10^, Figure S7A) and mice (r = 0.864, p < 10^-10^, Figure S7B). This correlation holds true for cycling genes (r = 0.641 in yeast and r = 0.836 in mouse, Figures S4C and S4D) and cycling genes show higher selective pressure on energetic cost (p < 7.03×10^-139^ in yeast and mouse, Figures S4E and S4F). These results imply that highly-expressed genes, together with cycling genes, are under strong selection against energy expenditure.

### Both Circadian Expression Peaks are Under Cost Selection

Circadian genes often cluster into two peak phases, with one peak in the first half of oscillation and the other in the second half of oscillation (the early peak and late peak, respectively). In yeast the number of genes located in the early peak is larger than that of genes in the late peak (1,708 genes versus 2,863 genes, Figure 5D), this number is larger in the late peak in mice (2,325 genes versus 2,832 genes, Figure 5D). We examined whether the two peaks are both under selection for reducing energetic cost and if there are any differences in selective pressure between them. In yeast,1,637 out of 1,708 genes (95.84%) are under selection during early-peak expression, while 2,778 out of 2,863 genes (97.03%) are under selection in the late peak. During the early peak of mice, 190 (8.17%) genes are under selection 122 (4.31%) in the late peak. For the cycling transcriptomes of both species, a significant enrichment of cost-selected genes and cycling genes can be detected in both peaks (OR > 3 and p < 7.61×10^-16^ in both comparisons). This shows that selective pressure has little bias for one peak over the other. We then asked whether the cost of a group of genes can reach the selection threshold for both peaks in mice. We found that the co-expression of at most 11 genes can pass the threshold for being selected against in both peaks, with 95% confidence (Figure 5C). This is far fewer than the number of phase-aligned genes in each peak (191 genes on average). The above results suggest the two peaks of rhythmic gene expression are important elements for removing unnecessary transcripts.

### Model: Energetic Cost Imposes a Dominant Constraint on Cyclical Expression

The common patterns of cycling gene distribution we observed, together with the cost analysis of cycling genes, lead us to propose the energetic cost optimization model to explain the evolutionary behavior of cycling genes. According to this model (Figure 6), the function of transcripts can be roughly classified into “always needed” and “temporally needed”, both of which are condition- and tissue-dependent. For the genes that are highly expressed, reducing their energetic cost is always required. If highly-expressed genes are always needed, they will rarely oscillate. However, if those highly-expressed genes are temporally needed, their oscillatory behavior will be heavily regulated. On the other hand, lowly-expressed genes experience less selective pressure on energetic cost regardless of whether they are temporally needed or not. In summary, the selection constraints of energetic cost on highly-expressed genes with temporally-controlled expression explains the major architecture of the cycling transcriptome.

**Figure 6.**
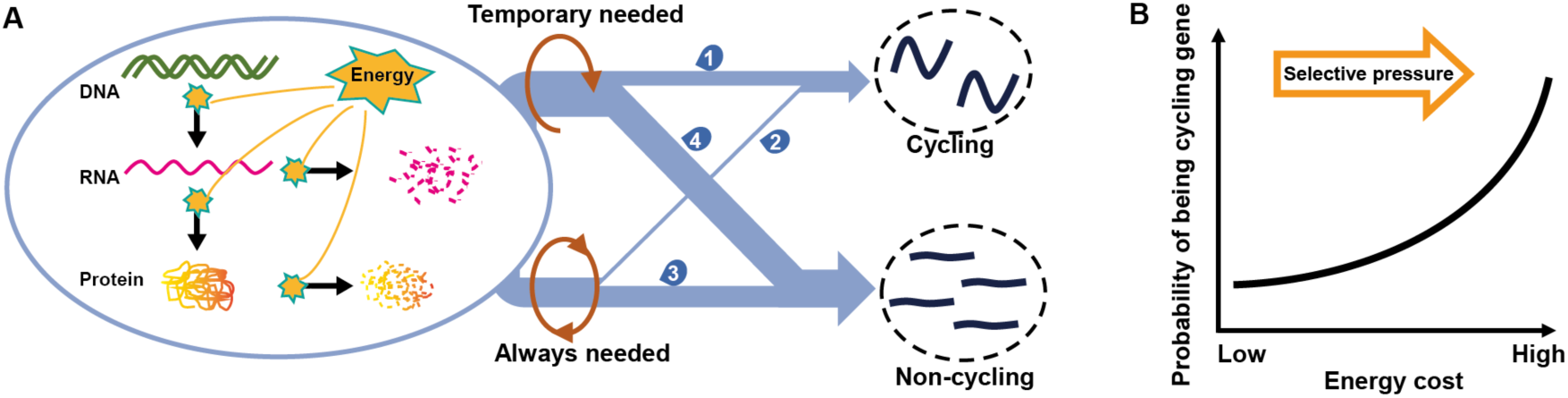
Modeling the cycling transcriptome via energetic cost optimization. (A) The amount of energy consumed during gene expression activities (transcription, translation, RNA synthesis and degradation, protein synthesis and degradation, etc.) is potentially subject to natural selection. The selection strength experienced by a gene depends on its total energetic costs, which is largely determined by its expression level. Expression of genes whose functions are temporarily needed or related to tissue-specific processes are more likely to undergo rhythmic regulation (1). Expression of genes whose function is constitutively needed or related to basic cellular processes are less likely to undergo rhythmic regulation (2). Non-cycling genes are observed in most of the cases as they can originate from either constitutively-needed genes (3) or temporarily-needed genes with an energetic cost that is too low to pass the selection threshold (4). (B) The relationship between energetic cost and cycling gene identity. The probability of being a cycling gene increases as selective pressure against energetic cost increases. Lowly-expressed genes typically consume less ATP during their expression and thus are under less selective pressure to be rhythmically regulated.

### Computational Simulation of the Energetic Cost Constraint Reproduces the Conserved Patterns of Cycling Genes

Finally, by performing a large-scale computational simulation, we tested whether our model of selection against energic cost can reproduce the conserved patterns observed for cycling genes. We randomly generated 10,000 protein-coding genes, each of which has a sequence length of 300 amino acids. These genes were then assigned to a functional matrix containing 1,000 functions, in which the functional distribution across time is followed by a normal distribution The genes were assigned to the function matrix based on a negative linear relationship so that most genes participate in just a few functional terms. For each function, the expression requirement of these genes was designated with a random number between 10-100. Thus, the required expression level for a specific gene at a certain time was determined by considering all the function terms in which this gene participated. For each gene expressed, a stochastic mRNA expression profile based on this required expression level was generated at each cycling time point. Each mRNA molecule was assumed to be translated 100 times in the simulation. For each coding sequence, we calculated the total energetic cost of its expression, which was determined by multiplying the expression level with the cost of a single mRNA/protein molecule (assuming other cost during transcription and translation as a constant *c*). We assumed differences between the cost of actual expression and required expression (Δcost) to be under natural selection. Selective strength was set to be positively correlated with Δcost. Finally, the expression level for each simulated gene was reevaluated based on the reduced energetic cost after selection and cycling genes were identified.

As shown in the bottom panel of Figure 3, the final identified cycling genes exhibited higher expression abundance and larger relative amplitude than non-cycling genes in the simulation. In addition, the proportion of cycling genes increases gradually as the expression level or rAMP increases. Thus, all four common distribution patterns of cycling genes can be reproduced. On the other hand, simulated data without selective constraint on cost does not exhibit the observed patterns (Figure S8). Our large-scale simulation also predicts that the number of cycling genes detected should increase as the selective strength (*s*) increases (Figure S9). If there is little selective pressure (*s* = 0.01), only 324 genes are detected to be rhythmically expressed in the simulation. However, if the selective strength is large (*s* > 0.11), more than 5000 genes can be detected as oscillating genes. The effect of selective strength on the number of cycling genes is consistent with observations that less oscillating genes are detected in aged tissues compared with adult tissues^21,47^.

## Discussion

Although the individual components of the molecular circadian clock are not conserved across species, the autoregulatory negative feedback loop they form is conserved. Likewise, the output of the regulatory network varies greatly between different species and/or environmental conditions, but whether or not these transcriptomes exhibit consistent design patterns is unknown. By comparing 14 diverse cycling transcriptomes in eight species, including human, mouse, fruit fly, Arabidopsis, yeast, *Neurospora crassa*, green algae and cyanobacteria, we discovered four conserved patterns of cycling genes. Cycling genes were highly expressed and displayed large regulation ranges in comparison to non-cycling genes. Additionally, the proportion of cycling genes increased with both mean expression and relative regulation range. Moreover, our calculations showed that cycling genes can be correctly predicted based on their mean expression level and relative range of expression.

In order to explain the conserved distribution patterns of rhythmic genes, we tested the hypothesis that cycling behavior can effectively optimize overall transcriptional energy expenditure. In species with a large effective population size such as yeast, the oscillation of more than 95% of genes was directly under this selection. In species with a relatively small effective population such as mice, the co-oscillation of as few as 9 genes was subject to this selection at a 95% confidence level. In addition, both the early and late peaks of rhythmic expression are under this selection. These results suggest that rhythmic gene expression can effectively remove unwanted transcripts by selecting against their energetic cost, especially for those genes that are highly expressed. In addition, the importance of phase alignment of gene expression is highlighted in species with a relatively small effective population size (such as mammals) in which the co-expression of phase-aligned transcripts experiences selection.

Our computational model based on constraining energetic cost completely reproduces the four conserved distribution patterns of rhythmic genes. Our model predicted that the number of rhythmically expressed genes increases with selection pressure on energetic cost, which is consistent with observations that young tissues generally have more oscillating genes than aged tissues^21, 47^. This suggests that circadian disruption in aged tissues follows the general evolutionary principle that selection pressure in older individuals is significantly decreased^48, 49^. Still, the aging process may involve distinct transcriptional networks in different tissues^21, 47, 50^.

The core circadian clock is generally considered to have at least two origins^6,51^, with convergent evolution promoting its origin and also the formation of the core design motif (the negative feedback loop). Convergent evolution has thus independently allowed molecular adaptation to the ∼24-hour rotation of the Earth in all species. Our findings indicate that the transcriptional output of this network has also obtained a consistent design principle, which suggests that convergent evolution might have also shaped this aspect of circadian biology. However, to what extend convergent evolution affected circadian output requires additional work.

Noncoding RNAs (ncRNAs) are pervasively transcribed in the genome, with many being regulated in a circadian manner^18, 52, 53^. As noncoding RNAs are typically expressed at low levels and not translated into proteins, our results indicate that they should generally experience less selection during daily oscillation. This also suggests that ncRNAs should exhibit less oscillatory behavior than mRNAs. This is consistent with the observation that ncRNAs are less likely to oscillate and no consistent ncRNAs are found in more than five mouse tissues^18^. Our model suggests that there is a smaller proportion of cycling ncRNA in aged tissues compared to young tissues, which can be tested with future experimentation.

Our work shows that circadian regulation of highly-expressed genes is an effective way to optimize the energetic cost of the transcriptome. Highly-expressed genes also contain a lower numbers of introns, which is energetically favorable for transcription^54^. On average, highly-expressed genes contain shorter 3’ UTRs and encode smaller proteins than other genes in the genome^55, 56^. Cycling-based regulation therefore appears to be a more flexible approach for directly affecting expression than interrupting the coding regions with introns. The liver is synchronized with diurnal-based signals, including feeding-fasting and light-dark conditions, situations in which liver mass and cell size are rhythmic^57^. The importance and universal significance of energetic cost optimization on the design of circadian architecture thus deserves more quantitative and systematic investigations, especially as it relates to metabolic and homeostatic processes.

## Methods

### Sources for cycling transcriptomes used in comparative analysis

Both microarray and RNA-sequencing cycling datasets were collected for eight species: human: 2 datasets; mouse: 2 datasets; *Drosophila melanogaster*: 2 datasets; yeast: 2 datasets; *Neurospora crassa*: 1 dataset; *Arabidopsis thaliana*: 3 datasets; *Chlamydomonas reinhardtii* (green algae): 1 dataset; and *Synechococus elongates*: 1 dataset. For human, circadian measurements on microarray data from both blood and skin were obtained (GSE39445 and GSE112660)^26, 27^. For mouse, circadian data for the liver and kidney (light/dark condition) were filtered for inclusion in our analysis, as these two tissues represented the organs with the highest numbers of cycling genes (GSE54650)^18^. For *Drosophila melanogaster*, RNA-sequencing data from both young and old fly heads were downloaded, and mitochondrial proteins were excluded from downstream analysis as they are overly abundant in those samples (GSE81100)^28^. The RNA-sequencing *Neurospora crassa* dataset^30^, which was profiled from a constant-dark condition, the RNA-sequencing data from green algae (GSE62671)^32^, the microarray data from cyanobacteria (GSE14225)^33^, and yeast metabolic cycling datasets under high- and low-glucose conditions (GSE57683)^29^ were downloaded and processed. Finally, for Arabidopsis, we obtained circadian data from three tissues: whole leaf, mesophyll (M), and vacuum (VA) under short day conditions (8 hours light and 16 hours dark) for comparative analysis (GSE50438)^31^. For microarray datasets, probes which mapped to the same locus were averaged to avoid duplicated gene identifiers. Finally, protein coding genes that have expression profiles in greater than half of all samples in each cycling transcriptome were considered as expressed genes for the downstream analysis (193,492 transcripts).

### Identification of cycling genes in each species

Gene expression measurements from all cycling transcriptomes were processed with the same workflow for identifying cycling transcripts (Data Table S2). To confidently identify rhythmically-expressed genes, the meta2d function from MetaCycle^34^ was utilized to calculate the period, phase, and amplitude for each cycling transcript, with parameter setting cycMethod = c("ARS", "JTK", "LS"). The transcripts with q values < 0.05 (under Benjamini-Hochberg procedure) were considered significantly cycling transcripts.

### Computational determination of orthologous genes

We downloaded the protein sequences of 8 species from the ENSEMBL database (http://asia.ensembl.org/). For each genome, coding genes with the longest protein sequences of that region were extracted and then OrthoFinder (v2.2.7)^58, 59^ was run to build the orthologous gene groups for each species pair (Data Table S5)). All one-to-many and many-to-many homologous gene pairs were retained in the analysis and then “Diamond gene sequence alignment” was used for sequence alignment and “fasttree” for phylogenetic tree building, with all other parameters set as default. In the comparative analysis of orthologous genes in different tissues of species pairs, results from average values of pairwise comparisons between them were reported.

### Functional enrichment analysis on cycling genes

Functional analysis of cycling genes was performed by utilizing the “Gene Ontology Resource” website (http://geneontology.org/)^35,36^. The cycling genes in each cycling transcriptome were used as an input gene list while the background genes were set as all expressed genes in the given transcriptome. Then the “PANTHER” classification system was chosen for the downstream enrichment analysis^60^. In the "Annotation Data Set" option, "GO biological process complete" was selected, and for the "Test Type", "Binomial" option was selected to perform the “PANTHER” overrepresentation Test^37^. Finally, p values less than 0.01 were used to call significantly enriched biological process^37^.

### Displaying the expression of cycling genes in each species

In order to show the rhythmicity of cycling gene signals, the "pheatmap" package was used to display rhythmic expression patterns. Each row represents the cycling time point while each column represents cycling genes (q < 0.05, Benjamini-Hochberg procedure). Cycling genes were ranked by the "phase" of their oscillation in descending order.

### Comparative analysis of cycling genes and their enriched pathways

In the comparative analysis of overlapping cycling genes and functional pathways across species, pair-wise comparisons were used. For each comparison, the overlapping cycling genes or pathways were filtered out and odd ratios were calculated according to the enrichment significance, with an applied Fisher’s exact test. Then evolutionary distances among the 8 species were calculated by “OrthoFinder” and were integrated in the correlative analysis with the enrichment significance of cycling genes or their related pathways.

### Simulation on the effect of expression level in detecting cycling genes

Simphony^61^, an R package that can simulate data for both cycling and non-cycling genes, was used to simulate large-scale cycling expression. A total of 10,000 genes were simulated at 2-hour intervals and sampled over a two-day period. We defined 5,000 genes as highly expressed and another 5,000 as low-expressed by setting the baseline expression, with a five-fold difference in the mean value of these two groups. 2,500 of the 10,000 genes were defined as cycling genes. The parameter that distinguishes cycling and non-cycling genes was their amplitude, which was defined as 0 for non-cycling genes, 1 for low-expressed cycling genes, and 5 for high-expressed cycling genes. The phases of the cycling genes were set evenly distributed across different time points. We assumed that the expression data conform to the negative binomial distribution, in consistent with the RNA-seq data. In the subsequent simulation, we added 20% and 50% of “expression noise” to the simulated data, with the mean expression level of each gene remains the same.

### Prediction of cycling genes based on mean expression levels and expression ranges

To predict oscillating genes in each transcriptome, all the expressed genes were first ranked by their expression range and the top 95% highly-ranked genes were extracted and included in the subsequent simulation. Then the probability of a gene being predicted to be a cycling gene was based on its relative expression, as shown in the formula:

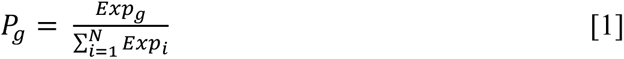

Where *P* represents the possibility of a certain gene to be selected as a cycling gene, *N* represents the number of selected genes in each condition and *Exp* represents the expression level in the simulation. In the control experiment, the probability of each gene predicted to be a cycling gene is not related to its expression level, *i.e.* all genes are equally selected. The total number of rhythmic genes was generated using the above simulation and compared with the actual data for each species.

### Calculation of energetic cost

The estimation of energetic cost for each cycling transcript in yeast and mice was largely based on previously published methodologies^41, 42^, with only slightly modifications which makes it more suitable for the situation of rhythmic expression. In brief, the calculation of the total cost of the transcription and translation of a gene in yeast and mice was based on the formula (3b) and (4) of the previous report^42^. For the cycling transcriptome, we assumed that half of nucleotides and amino acids are recycled for anabolic purposes and do not need to be synthesized *de novo*. To calculate degradation rates for each molecule, the experimentally measured mRNA half-lives and protein half-lives for the two species were collected from previously reported datasets^62–64^.

The cost of mRNA biogenesis was calculated by:

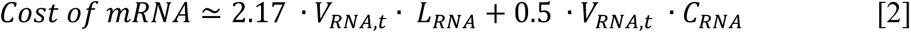

Where *V_RNA,t_* represents the mRNA synthesis rate for each gene at time point *t,L_RNA_* represents the length of a specific mRNA, and *C_RNA_* represents the energetic cost for transcribing a single mRNA molecule.

Then the cost of protein biogenesis was calculated by:

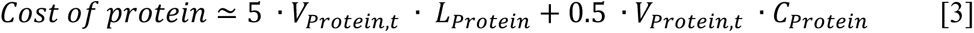

Where *V_protein,t_* represents the protein synthesis rate for each gene at time point *t,L_Protein_* represents the length of a specific protein sequence, and *C_Protein_* represents the energetic cost for translating a single protein molecule.

Then the mRNA synthesis rate was determined by the following formula:

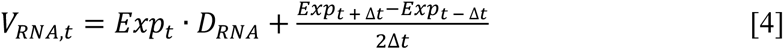

Where *Exp_t_* represents the expression level of one gene in a specific time point *t*, and *D_RNA_* represents degradation rate, which is calculated by ln2 divided by the half-life of the mRNA.

In mice, ribosome-profiling data^45^ were explored to estimate the corresponding synthesis rate for each circadian time point. We estimated the cycling protein synthesis rate by combining both the cycling transcriptome and ribosome profiling data from mouse liver. As the elongation rate is on average 6.8 amino acids per second in the mouse liver^65^, 6.8×3×3600=73.44k nucleotides can be translated in one hour. We estimate the protein synthesis rate of a certain mRNA (RPKM) in mouse liver by the following formula:

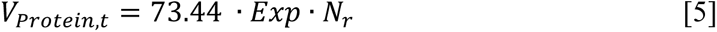

where *V_Protein,t_* represents the synthetic rate of mouse protein, *Exp* represents RPKM values of mRNA, and *Nr* is the number of ribosomes *per* 1000 bp.

In yeast we estimated cycling protein abundance first based on the ratio between protein copy number and mRNA copy number^66, 67^. We then calculated the protein synthesis rate with a formula similar with that of the mRNA synthesis rate (formula 2 of this manuscript above). For the genes that lack experimentally-measured parameters, the median values of all genes were used in the calculation. Finally, the fractional energetic cost per time point was computed as the cost of a particular gene divided by the total cost of all expressed genes at a given time point. If this value (fractional energetic cost) is larger than the selective threshold of the population, this gene is under selection for energetic cost.

### Computational simulation of cost constraints on cycling transcriptome

A computational model was developed to simulate the effects of energetic cost constraints on rhythmic gene expression. In the simulation, 10,000 genes were generated, each consisting of 300 amino acids. 1,000 functions were randomly assigned to these genes according to the rule that most genes participate in a limited number of functions. Then we assigned those functions to 24-hour temporal organization which is based on a roughly normal distribution, *i.e.* more functions were active around noon than other times.

Then for each function that was engaged at a particular time point, the expression level of the genes in that function were randomly assigned values from 10 to 100. Then the expression level of each gene was calculated for all the cycling time points (needed expression). As gene expression is essentially stochastic, we assumed that the measured expression level ranges from 0.9 *x* needed expression level to 1.5 *x* maximum needed expression level of all cycling time points. Our central hypothesis is that cost selection exists for the transcript that are highly expressed. We assumed that for genes with an extra cost in their expression (Δcost), selection pressure scales with cost. We use *R* as the constraint ratio, which was determined by:

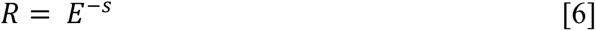

where *s* represents the selective strength, which is between 0 and 1, while *E* represents Δcost.

Finally, we calculated adjusted expression levels for each gene based on their constraint ratios, identified cycling genes based on their adjusted expression levels, and then characterized their distributions.

### Statistical significance tests

Statistical analysis was performed using R software (version 4.0.3). Wilcoxon rank sum tests were performed for the comparison of the conservation levels between and within species at the gene and/or pathway level. Pearson correlation tests were performed to examine correlations between averaged expression levels and cycling amplitudes and Fisher’s exact tests were used to examine the enrichments between cycling genes and the genes undergoing energetic cost selection. p < 0.05 were considered significant in the analysis and exact p values were reported. q value < 0.05 from MetaCycle^34^ was used as the significance cutoff for the identification of cycling genes.

### Data and Software Availability

Datasets and the codes used to generate the results in this research have been deposited online in GitHub and are freely accessible to the public (https://github.com/WangLab-SINH/RhythmicExpressionConstraints).

## Author contributions

G.-Z.W. conceived and designed the study. Y.C., Y.C., and L.S. conducted the analysis. Y.C., Y.C. and G.-Z.W. wrote the manuscript. All authors contributed to the Discussion.

## Acknowledgement

We thank Daniel J. Araujo, Chris Turck, and Mindian Li for helpful discussion and critical reading of the manuscript. This work is funded by the National Natural Science Foundation of China (Nos. 81827901, 31600960 and No. 31871333), Shanghai Municipal Science and Technology Major Project (Grant No. 2017SHZDZX01).

## Supplementary Information

**Figure S1.**
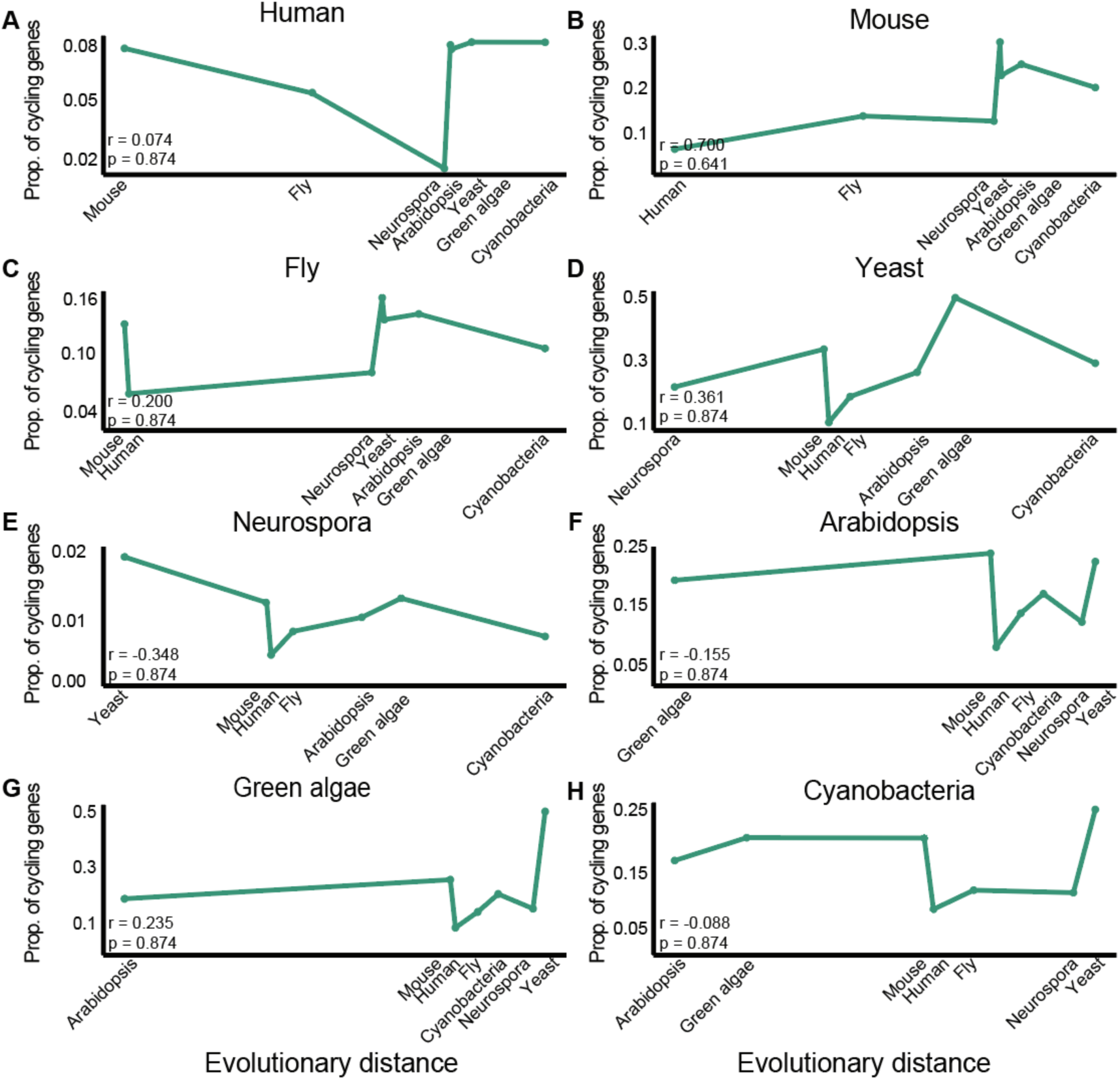
Relationships between shared cycling genes and evolutionary distance. (A-H) Line charts showing relationships between proportions of cycling genes and evolutionary distances amongst species considered. The horizontal axis represents the evolutionary distance, while the vertical axis represents the proportion of cycling genes, which is defined by the Jaccard coefficient of cycling genes between the two species. Pearson correlations were used to calculate relationships between evolutionary distances and Jaccard coefficients, with p > 0.6 used for all comparisons. p was corrected using a Benjamini-Hochberg procedure. Similar results were obtained when an odds ratio was used instead of the Jaccard coefficient.

**Figure S2.**
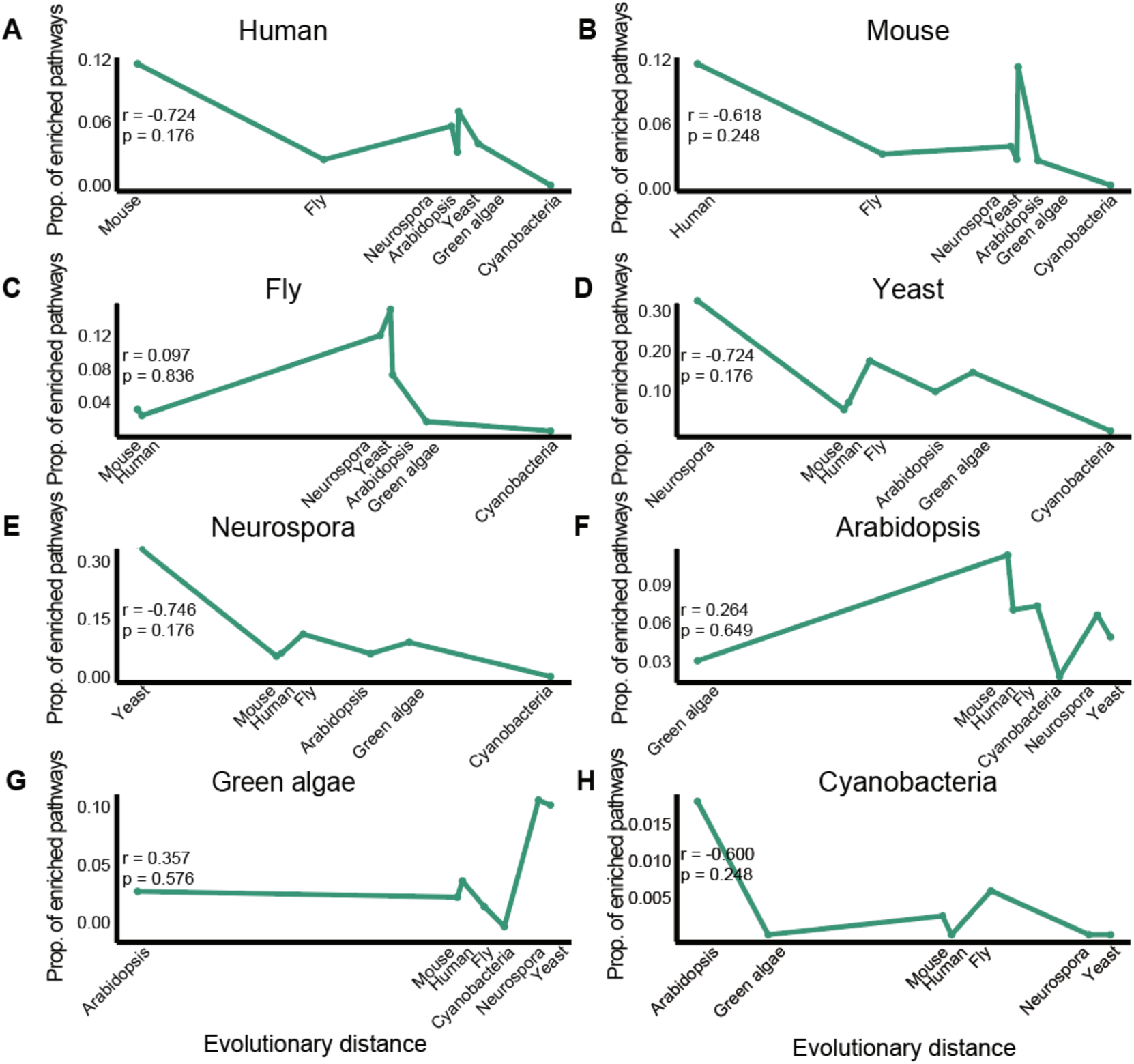
Relationships between conserved enriched pathways of cycling genes and the evolutionary distance. (A-H) Line charts displaying the relationship between the proportion of enriched biological pathways for cycling genes and the evolutionary distance between species. The horizontal axis represents the evolutionary distance, and the vertical axis represents the Jaccard coefficient of the circulating genes between the two species. Pearson correlation was used to calculate the correlation between evolutionary distance and the Jaccard coefficient. p values were corrected using a Benjamini-Hochberg procedure. Similar results were obtained when we applied an odds ratio instead of the Jaccard coefficient.

**Figure S3.**
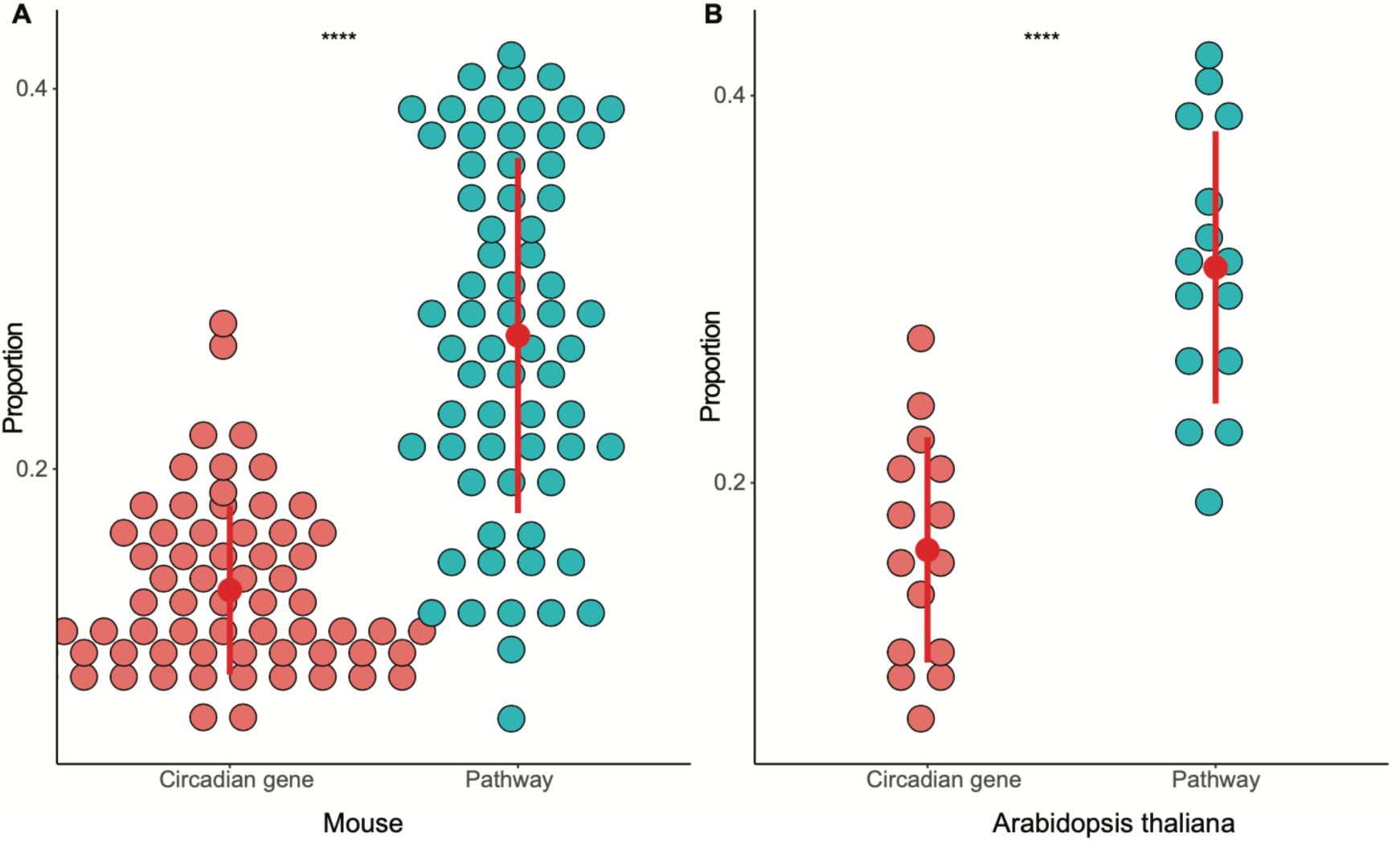
Comparisons between the conservation of circadian genes and their corresponding enriched pathways. Boxplot showing the proportion of shared circadian genes compared with the proportion of their shared enriched pathways for the 12 mouse (A) and the 6 Arabidopsis (B) tissues considered. Red dots represent the proportion of shared circadian genes, and green dots represent the proportion of shared enriched pathways. Wilcoxon rank sum tests were used to compare the differences between the two groups and **** indicates p < 10^-5^.

**Figure S4.**
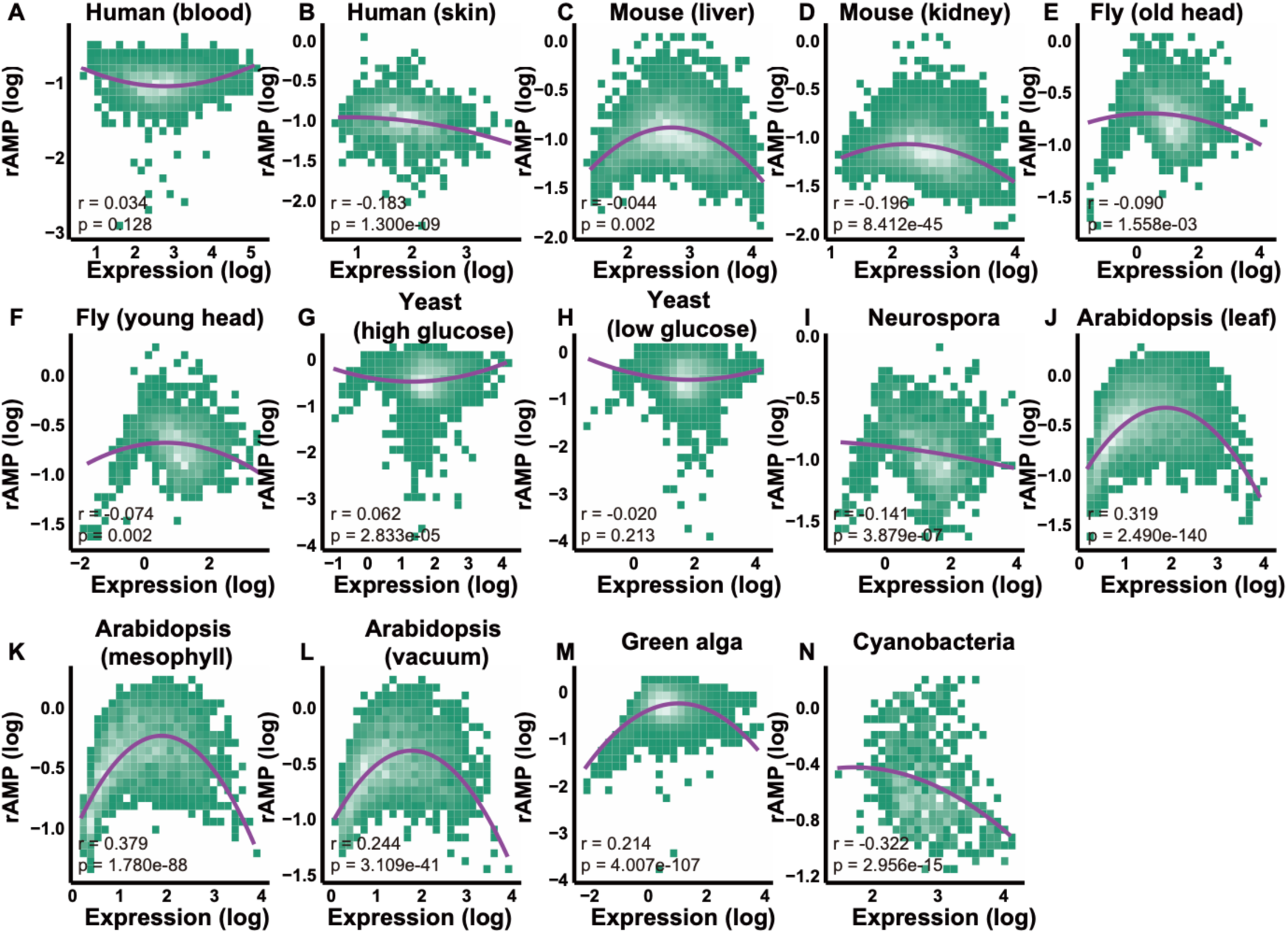
The relationship between rAMP and mean expression. (A-N) Scatter plot representing the relationship between expression level and rAMP, where x-asix shows the expression level, and y-asix shows the rAMP value. Both of x-asix and y-asix were logarithmic transformed.

**Figure S5.**
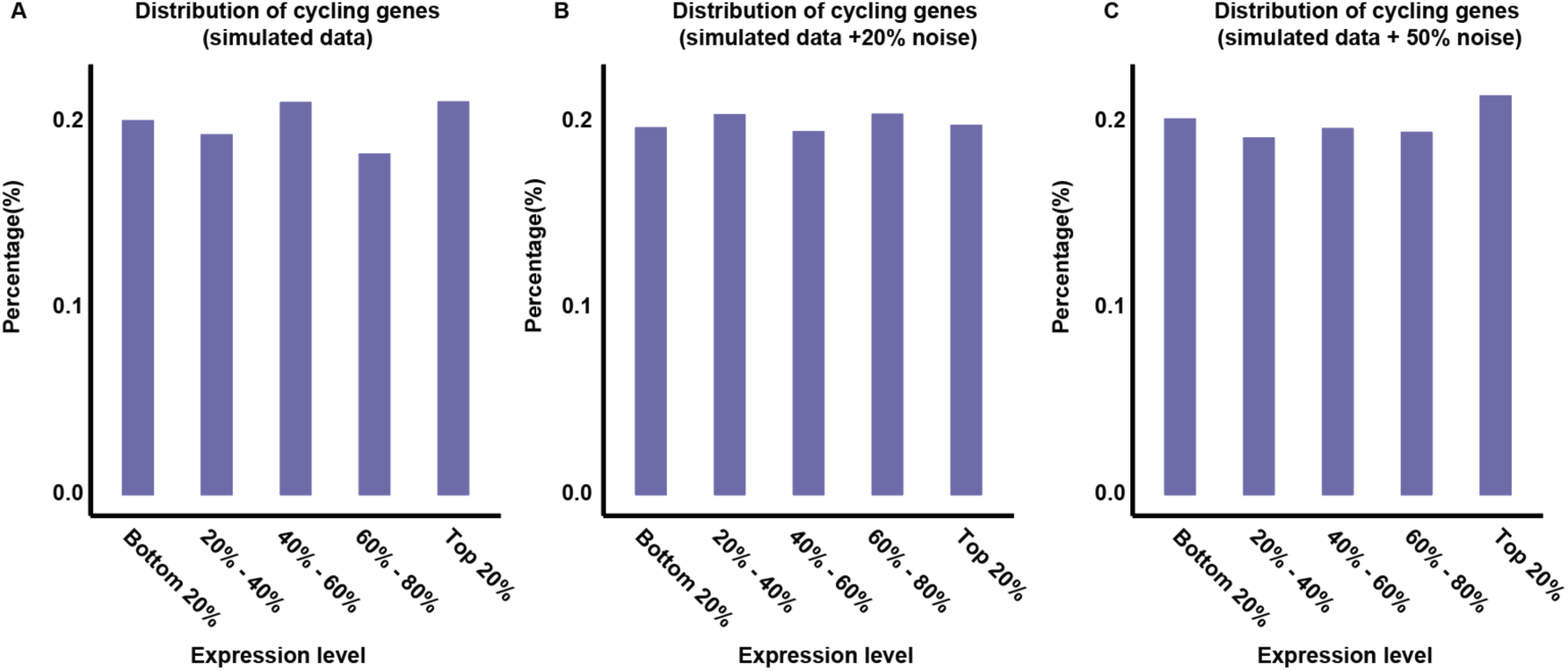
Relationship between noise and cycling gene detection. (A) Proportions of cycling genes grouped by mean expression in 10,000 simulated genes. (B) Proportions of cycling genes grouped by mean expression in simulated data with 20% noise added. (C) Proportions of cycling genes grouped by mean expression level in simulated data with 50% noise added.

**Figure S6.**
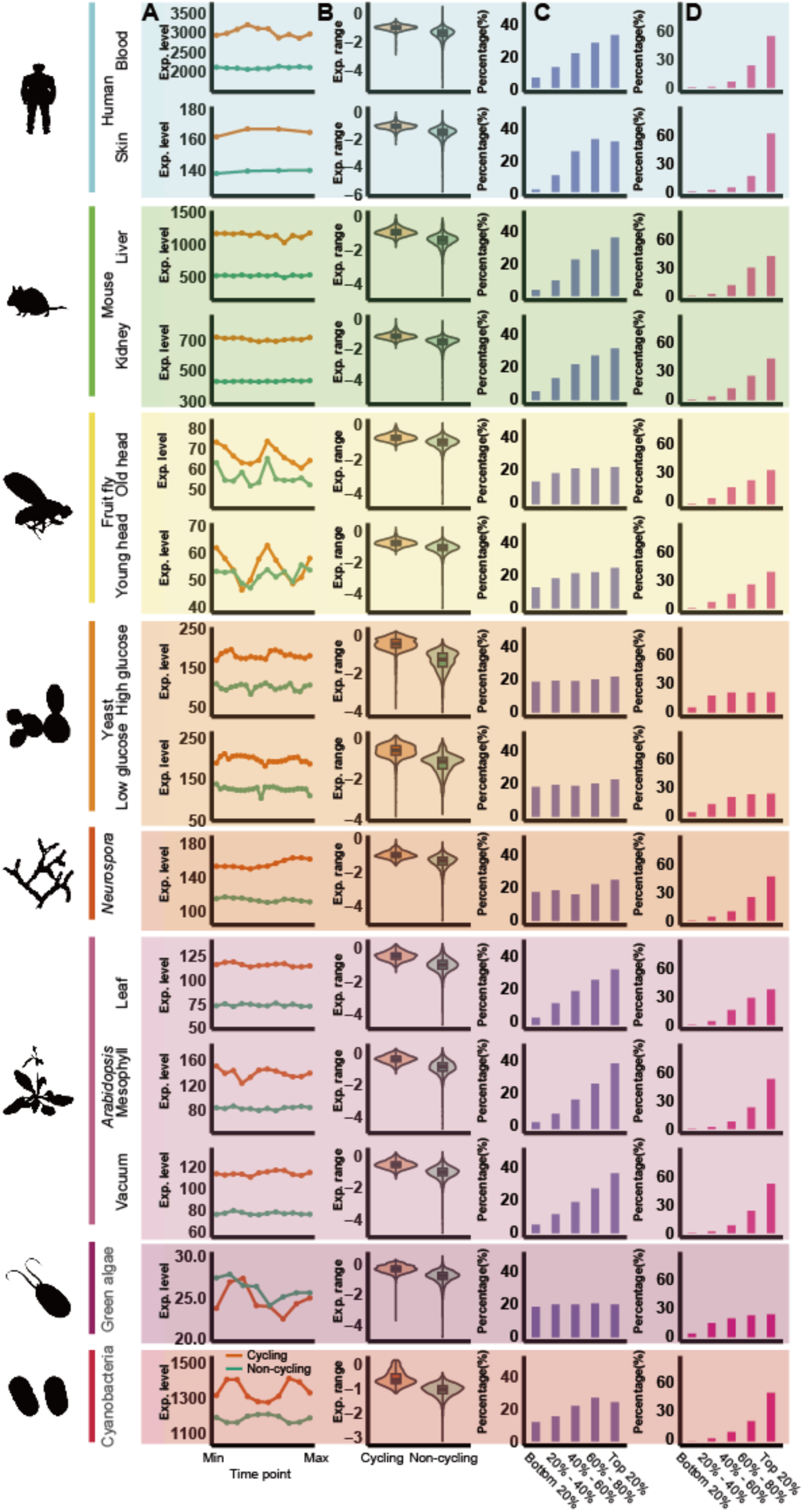
Four patterns remained after removing the 20% lowly expressed genes in each cycling transcriptome. (A) Mean expression levels of cycling genes versus other genes for each time point. Red lines represent cycling genes while green lines represent non-cycling genes in each plot. (B) Violin plots showing comparisons of rAMP between cycling genes and other non-cycling genes. Red boxes represent cycling genes while green boxes represent non-cycling genes. (C-D) Proportions of cycling genes in each category grouped by mean expression level (C) and rAMP (D). Percentages were calculated as the number of cycling genes in that group divided by the total number of cycling genes identified in the cycling transcriptome of each species.

**Figure S7.**
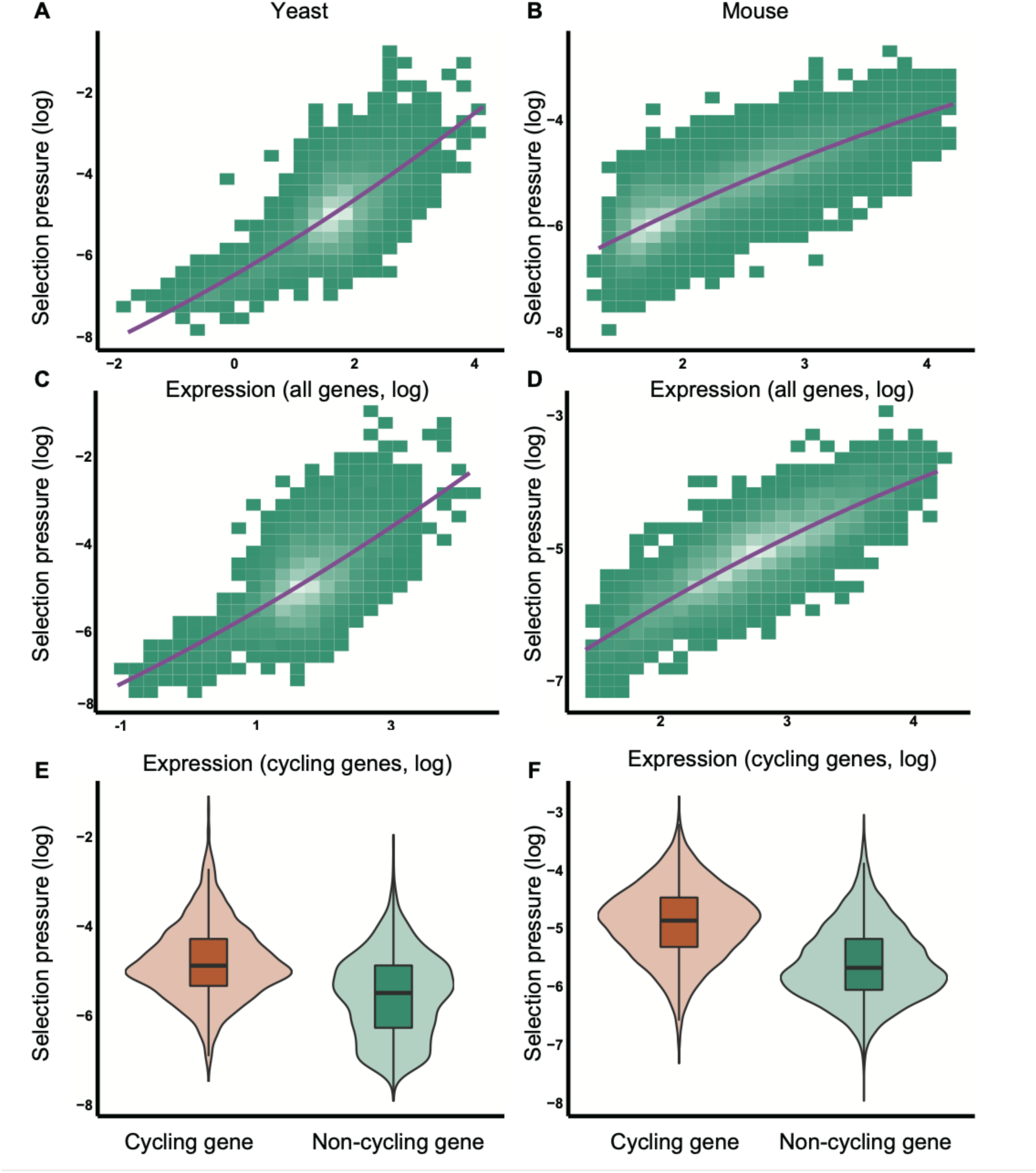
Relationships between selective pressure and gene expression level. (A-B) Scatter plot representing the relationship between expression levels and selection pressure in yeast (A) and mice (B). (C-D) Scatter plot representing the relationship between the expression levels of cycling genes and their corresponding selection pressures in yeast (C) and mice (D). (E-F) Violin plot showing a comparison of the selection pressures on cycling and non-cycling genes in yeast (E) and mice (F). Orange group represents cycling genes and green group represents non-cycling genes.

**Figure S8.**
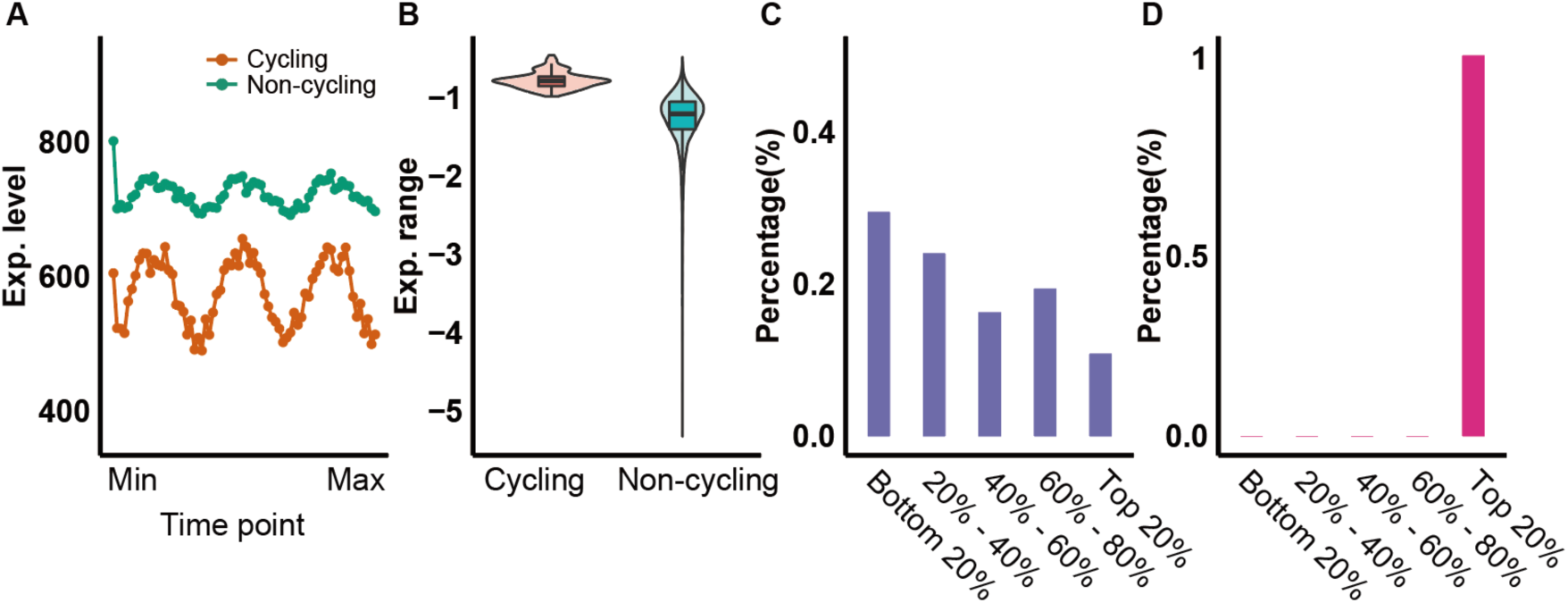
The simulated data without selection pressure did not exhibit the four observed patterns. (A) Mean expression levels of cycling genes versus other genes for each time point. Red lines represent cycling genes while green lines represent non-cycling genes in each plot. (B) Violin plots showing comparisons of rAMP between cycling genes and other non-cycling genes. Red boxes represent cycling genes while green boxes represent non-cycling genes. (C-D) Proportions of cycling genes in each category grouped by mean expression level (C) and rAMP (D). Five groups were calculated, from the bottom 20% to the top 20%. Percentages were calculated as the number of cycling genes in that group divided by the total number of cycling genes identified in the cycling transcriptome of each species.

**Figure S9.**
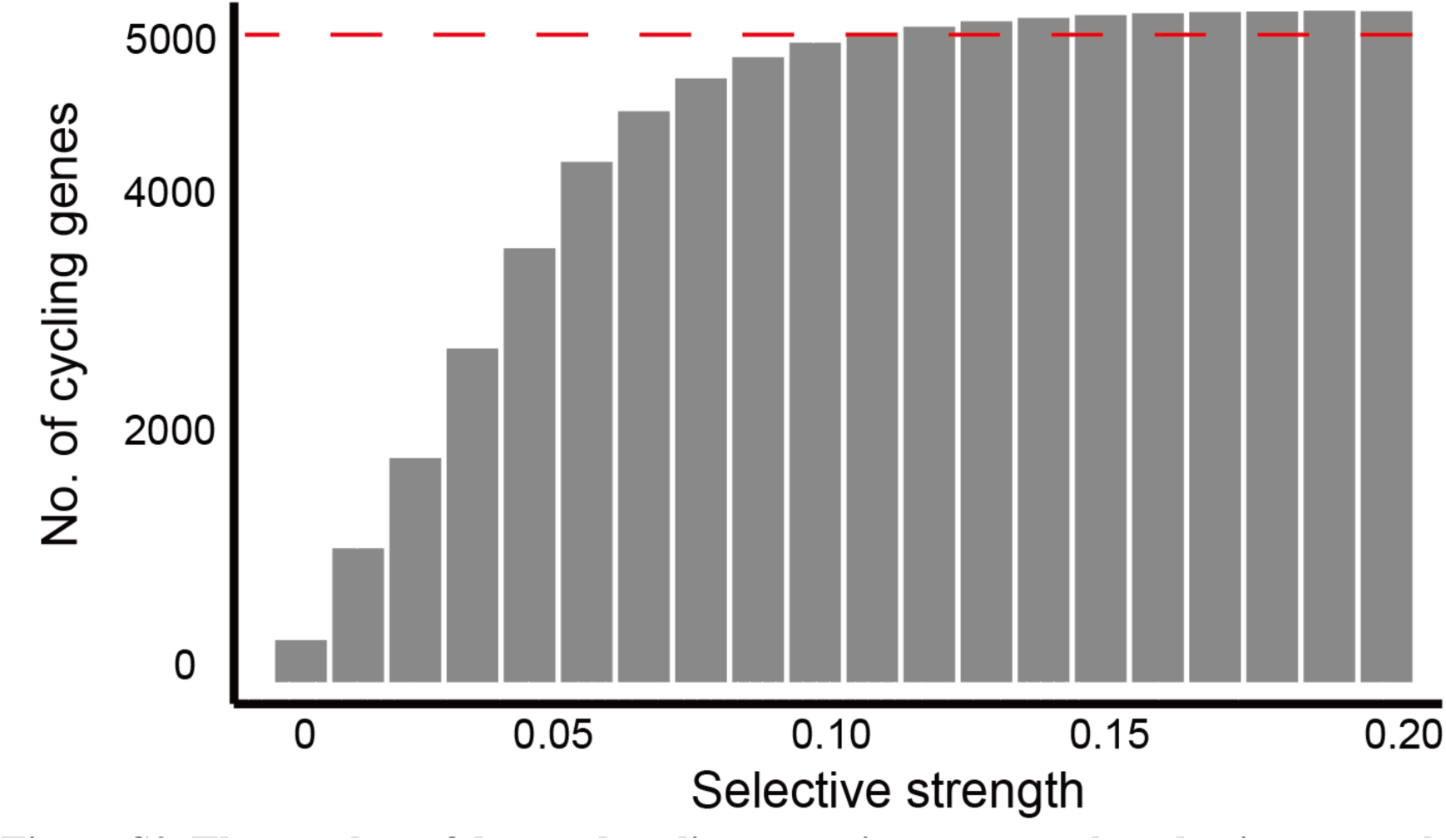
The number of detected cycling genes increases as the selective strength on energetic cost increases. x-axis represents selective strength while y-axis represents the number of cycling genes.

**Table S1. Information on the cycling transcriptome in 8 species** (Related to Figures 1).

Provided as separate .doc file

**Table S2. Information on the identified cycling genes in 8 species (Related to Figures 1 and 3).**

Provided as separate .xls file

**Table S3. Enriched functional pathways for the cycling genes of 8 species (Related to Figure 2).**

Provided as separate .xls file

**Table S4. Fractional energetic cost of all genes in yeast and mouse (Related to Figure 5).**

Provided as separate .xls file

**Table S5. Orthologous gene groups in 8 species** (Related to Figure 1 and 2).

Provided as separate .xls file

